# Specificity of RNA folding and its association with evolutionarily adaptive mRNA secondary structure

**DOI:** 10.1101/441006

**Authors:** Gongwang Yu, Hanbing Zhu, Xiaoshu Chen, Jian-Rong Yang

**Affiliations:** Program in Cancer Research, Zhongshan School of Medicine, The Fifth Affiliated Hospital, Sun Yat-sen University, Guangzhou, China 510080; Department of Biology, Zhongshan School of Medicine, Sun Yat-sen University, Guangzhou, China 510080; Zhongshan School of Medicine, Sun Yat-sen University, Guangzhou, China 510080; Department of Medical Genetics, Zhongshan School of Medicine, Sun Yat-sen University, Guangzhou, China 510080; RNA Biomedical Institute, Sun Yat-Sen Memorial Hospital, Sun Yat-sen University, Guangzhou, China 510120

## Abstract

Secondary structure is a fundamental feature for both noncoding and messenger RNA. However, our understandings about the secondary structure of mRNA, especially for the coding regions, remain elusive, likely due to translation and the lack of RNA binding proteins that sustain the consensus structure, such as those bind to noncoding RNA. Indeed, mRNA has recently been found to bear pervasive alternative structures, whose overall evolutionary and functional significance remained untested. We hereby approached this problem by estimating folding specificity, the probability that a fragment of RNA folds back to the same partner once re-folded. We showed that folding specificity for mRNA is lower than noncoding RNA, and displays moderate evolutionary conservation between orthologs and between paralogs. More importantly, we found that specific rather than alternative folding is more likely evolutionarily adaptive, since it is more frequently associated with functionally important genes or sites within a gene. Additional analysis in combination with ribosome density suggests the capability of modulating ribosome movement as one potential functional advantage provided by specific folding. Our findings revealed a novel facet of RNA structome with important functional and evolutionary implications, and points to a potential way of disentangling mRNA secondary structures maintained by natural selection from molecular noise.

## Introduction

Single-stranded RNA molecules spontaneously fold into various secondary structures through intramolecular base pairing. These structures, especially the evolutionarily conserved ones, are considered essential for the function of noncoding RNAs such as tRNA, miRNA (Hasler and Meister 2016), snRNA (Dunn and Rader 2010), snoRNA (Washietl, et al. 2005) and rRNA (Petrov, et al. 2014). For the coding/messenger RNA, secondary structures are implicated in localization(Martin and Ephrussi 2009), (de-)stabilization (Hollams, et al. 2002; Meisner, et al. 2004) and editing(Tian, et al. 2011) of RNA molecule, as well as the regulation of translational repression (Kertesz, et al. 2007; Gu, et al. 2010), translational elongation speed (Shalgi, et al. 2013; Yang, et al. 2014; Burkhardt, et al. 2017) and co-translational protein folding (Richter and Coller 2015; Faure, et al. 2016). Furthermore, natural selection for functional RNA secondary structure constrains the evolution of RNA sequence (Park, et al. 2013; Li, et al. 2016; Li and Zhang 2018). Therefore, study of RNA structome is of fundamental importance in RNA biology and evolutionary biology. More recently, development of high throughput sequencing(HTS)-based assay for RNA secondary structure, such as PARS (Kertesz, et al. 2010), icSHAPE (Flynn, et al. 2016), FragSeq (Underwood, et al. 2010), structure-seq (Ding, et al. 2014), RPL (Ramani, et al. 2015), PARIS (Lu, et al. 2016) and SPLASH (Aw, et al. 2016) has started to reveal a more complete picture of the secondary structure of different RNA molecules.

Despite these progresses, our understandings about the secondary structure of mRNA, especially for the coding regions, are mostly anecdotal. One major obstacle is that mRNA molecule is frequently treaded into translating ribosomes, which can only accommodate single-stranded mRNA. This repeated disruption by translation triggers frequent re-folding of mRNA and presumably makes experimental detection of consensus structure difficult *in vivo*. Indeed, active unfolding of secondary structures is found for mRNAs in yeast, suggesting a minor role of thermodynamics for mRNA folding *in vivo* (Rouskin, et al. 2014). On the contrary, the majority of structured noncoding RNAs have one single functional secondary structure, which is usually stabilized by protein molecules.

In this context of frequent re-folding, the physical proximity of two linearly remote fragments in the same molecule by Watson-Crick base-pairing, or simply RNA “folding”, could be divided into two categories. On the one hand, a fragment of RNA that always pairs up with the same remote fragment once re-folded, has specific folding. On the other hand, RNA fragment capable of pairing with different remote fragments when re-folded, have nonspecific/alternative folding. A classic example of alternative folding is riboswitches, whose function relies on the exchange between two mutually exclusive conformations (Lemay, et al. 2006; Lipfert, et al. 2007; Lemay, et al. 2009; Whitford, et al. 2009).

Intuitively, mRNA folding should be less specific than noncoding RNAs due to the interference of translating ribosomes. For example, either counterpart in a particular folding could be excluded from any base-pairing due to ribosome occupation. The alternative folding thus formed likely retains until ribosome moves, because the kinetics of spontaneous exchange between alternative structures tend to be slow (Uhlenbeck 1995; Treiber and Williamson 1999; Woodson 2000). Additionally, frequent reorganization of local mRNA secondary structures by translating ribosomes gives mRNA ample opportunities to sample alternative (sub-)optima in the energy landscape, making mRNA more likely to have alternative folding compared to noncoding RNA. Indeed, it was recently found that about 20%-50% of top 50 mRNAs with the highest numbers of detected secondary structure have at least one pair of alternative structures, some evolutionarily conserved, suggesting that alternative structures are pervasive (Lu, et al. 2016).

Despite the unambiguous evidence for alternative folding in mRNA, as well as the importance of mRNA secondary structure, there is a lack of systemic investigation on the folding specificity of mRNA, let alone its functional and evolutionary impact. In particular, several questions revolving around folding specificity are critical to our understandings of RNA biology. For example, what is the average level of folding specificity of mRNA, and how does folding specificity affects the function of mRNA. Theoretically, if the structures of mRNA molecules from the same gene are too diverse, few of them can have significant function due to the negligible number of molecules assumed each structure. On the contrary, isolating foldings with strong specificity should help identify functional structures. From an evolutionary perspective, it is also important to find out whether alternative or specific folding is generally adaptive, and why is it so.

In search of answers to these questions, we studied the folding specificity of mRNA using high throughput sequencing data of RNA folding in yeast and mouse. We confirmed the reduction of folding specificity in messenger compared to noncoding RNA, and showed heterogeneity of folding specificity among genes and sites within the same gene. Furthermore, folding specificity, instead of diversity, is found more prevalent in important genes and sub-genic regions, suggesting that folding diversities are generally non-adaptive. Finally, by demonstrating the different ribosome stalling capacity of specific and non-specific folding, we provided one potential mechanistic explanation for the functional impact and evolutionary benefit imposed by specific RNA folding. Our results thus revealed a novel facet of RNA structome with important functional and evolutionary implications, opening new path towards better understandings of mRNA secondary structure.

## Results

### Estimation of folding specificity for mRNA

By definition, folding specificity of a RNA fragment can be estimated by its probability of pairing up with one or more remote fragments. Qualitatively, if two fragments pair up with 100% probability once folded, the folding is specific, or *vice versa*. Quantitatively, the level of folding specificity is determined by the number and relative frequency of alternative foldings, or in other words opposite of the diversity of folding partners. The experimental data informative for folding specificity had not been available until the recent advancement of HTS-based assays for RNA duplexes *in vivo* (Graveley 2016). In particular, RNA proximity ligation (RPL) followed by deep sequencing yield chimeric reads with ligation junctions in the vicinity of structurally proximate bases in yeast(Ramani, et al. 2015). In addition, psoralen analysis of RNA interactions and structures (PARIS) employed psoralen crosslinking to globally map RNA duplexes with near base-pair resolution in mouse cells(Lu, et al. 2016). With these datasets, a list of folding partners for each RNA fragment can be extracted and used to estimate folding specificity (See Materials and Methods). Note that both assays have yet to reach base-pair resolution, thus we hereby studied the specificity of folding instead of pairing.

To evaluate the diversity of folding based on the experimental data, we need a unified measurement taking into account both the number of alternative folding and the relative frequency of them. To this end, we borrowed the idea of Shannon’s metric of information entropy(Shannon 2001) (Shannon index), which quantifies the uncertainty in predicting the identity of a randomly chosen entity from a system. Shannon index is frequently used in ecology as an index for species diversity (the uncertainty of predicting the species randomly captured from a community) (Tuomisto 2010), or similarly for measuring molecular diversity in various biological systems (the uncertainty of predicting the sequence randomly picked from a pool of nucleic acid or protein) (Lin 1996; Medinger, et al. 2010; Chouari, et al. 2017; Mangin, et al. 2017), or RNA folding specificity as defined by the computationally predicted ensemble of secondary structures (Huynen, et al. 1997). Adapted from Shannon entropy, we measured the diversity of folding by *S*_*obs*_ = − Σ_*i,j*_*p*_*i,j*_ln*p*_*i,j*_. Here *i* and *j* are two bases on the RNA, and *p*_*i,j*_ is the probability that physical proximity between base *i* and *j* is observed among all the chimeric reads derived from the gene in RPL or PARIS assay (See Materials and Methods). To guard against the confounding effect of sequencing depth and number of relevant sites, *S*_*obs*_ is further compared to its theoretical maximal value 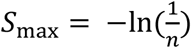, where *n* is the total number of pairs of nucleotides whose physical proximity are revealed by at least one chimeric read. Here *S*_*max*_ is essentially the information entropy when folding of all relevant bases are equally supported by the experimental data. The folding specificity is then defined as *S* = *S*_max_ − *S*_obs_, where higher *S* represents stronger folding specificity (**Fig. S1**). The formula for *S* is mathematically equivalent to Theil index, which is commonly used to measure economic inequality (Cowell 2003).

We first calculated the folding specificity of yeast mRNA using the RPL data(Ramani, et al. 2015). The recapitulation of folding specificity by *S* was confirmed by manual inspection of a couple of genes (**Fig. 1A**). For example, in YLR441C, none of the folding experimentally revealed is supported by more than one chimeric reads. Whereas for YPR154W, some foldings are supported by multiple chimeric reads with minor offset. The folding specificity of these two ORFs are respectively quantified as *S* = 0 and 0.12. Similarly, we estimated the folding specificity for all yeast mRNAs with at least 5 chimeric reads (**Fig.1B**). As a result, we found that the folding specificity of yeast mRNA varies greatly between genes: a substantial number of the genes show no measurable signal of folding specificity, whereas one gene (YDR420W) displays folding specificity comparable to that of noncoding RNA snR190 (**Fig. 1B**). The distribution of folding specificity of mRNA is similar when we only used yeast genes with 10 chimeric reads (**Fig. S2A**), or use folding specificity derived from mouse PARIS data (**Fig. S2B**). Additionally, the two biological replicates from mouse PARIS data allowed us to compare mRNA folding specificity of mRNA estimated by the two datasets, whose Pearson’s Correlation Coefficient is 0.48 (*P* < 10^−231^. **Fig. S2D**), suggesting that the heterogeneity of folding specificity is an intrinsic property of the transcriptome, instead of experimental noise.

**Figure 1.**
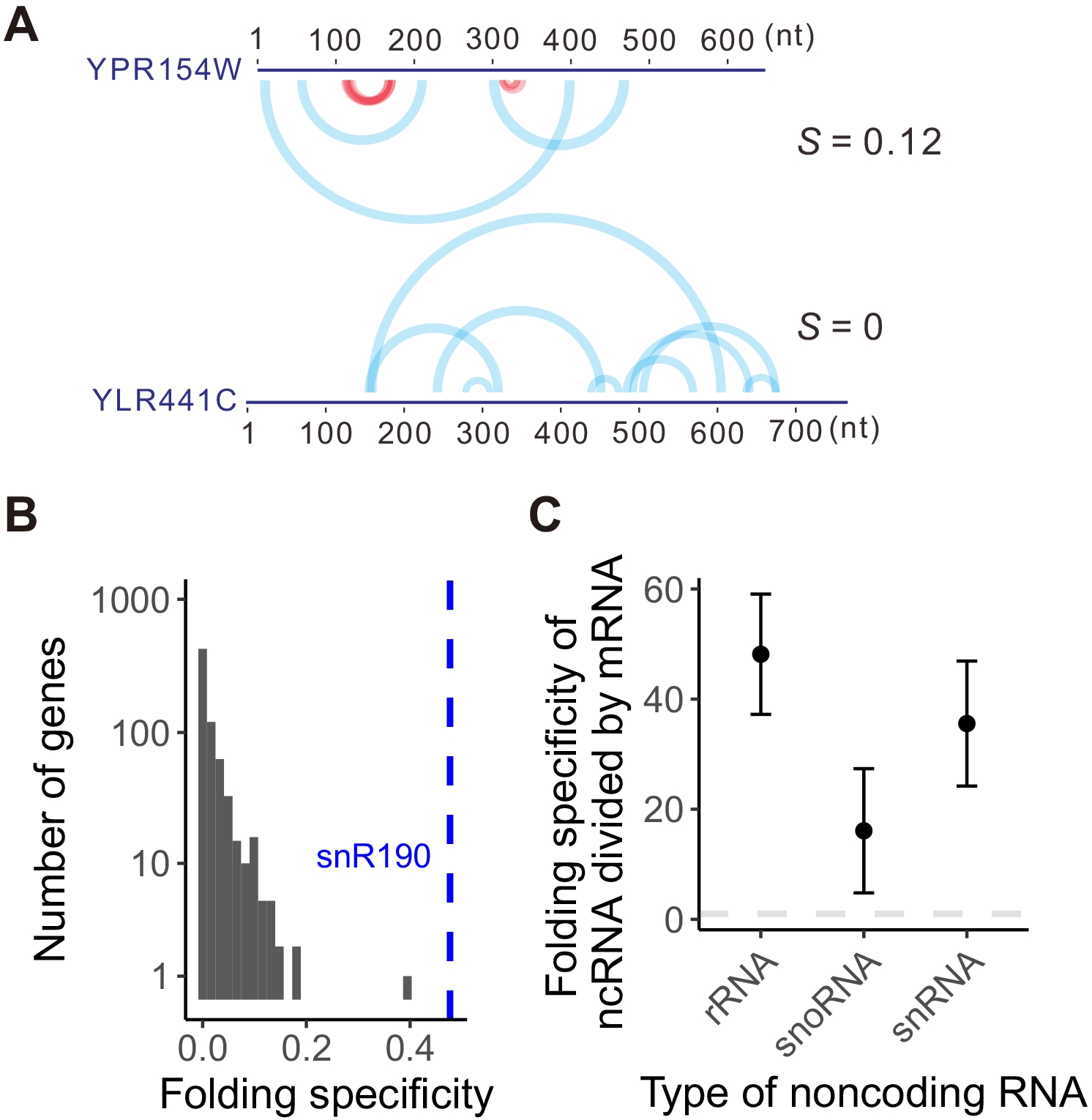
Folding specificity of yeast mRNA. (A) Two examples of folding specificity estimated from RPL data. Two yeast genes with their names, lengths and corresponding folding specificities (*S*) were shown. Each arch connects the two fragments that are folded together as suggested by one chimeric read in RPL data. The red arches support specific foldings with minor offset, whereas the blue arches support nonspecific foldings. The arches were transparent so that multiple chimeric reads supporting folding of the same pair of fragments will be shown as deeper color. Specific foldings are apparent for the top gene but absent for the bottom gene. The folding specificity is thus higher in the former (0.12) than in the latter (0). See also **Fig. S1**. (B) Distributions of folding specificity for mRNAs in yeast. A total of 697 mRNAs with at least 5 intramolecular chimeric reads were used. The folding specificity of a noncoding snoRNA snR190 is indicated by a dashed line, as a comparison. (C) Folding specificity is significantly stronger for noncoding RNA than mRNA in yeast. Major types of noncoding RNAs with folding specificity estimation were compared with that of mRNA. The ratio between the average folding specificity of the three types of noncoding RNAs and that of mRNA are significantly higher than 1 (the horizontal dashed line). Error bar indicates one standard error, estimated by bootstrapping the protein coding genes 1,000 times. Types of ncRNA presented include, ribosomal RNA (rRNA), small nucleolar RNA (snoRNA) and small nuclear RNA (snRNA).

We also compared the folding specificity of yeast mRNA with that of several types of noncoding RNA. It is well known that noncoding RNAs can fold into extensive secondary and tertiary structures, on which their functionality relies. Unlike mRNA, the secondary structure of noncoding RNA is not disrupted by translating ribosomes, but is instead usually stabilized by proteins. Therefore, the folding specificity of noncoding RNA is expected to be higher than that of mRNA. Indeed, the folding specificity of noncoding RNAs are significantly higher than that of mRNAs (**Fig.1C**). Similar results are observed for mouse PARIS data (**Fig. S2C**), reassuring the biological relevance of folding specificity. Note that *S* is usually small, which suggests weak signal for folding specificity, a phenomenon likely caused by both limited coverage of RPL/PARIS, and the frequent re-folding of mRNA *in vivo*.

### Folding specificity is an evolutionarily conserved molecular trait

To further assess the biological significance, we compared the folding specificity of one-to-one orthologs between yeast and mouse protein coding gene, and a moderate yet significant positive correlation was found between yeast and mouse (**Fig. 2A**). Considering the experimental noise and technical differences underlying the RNA folding data of yeast (RPL) and mouse (PARIS), the actual correlation should be stronger than currently revealed. Indeed, when we compared folding specificity of paralogous gene pairs in yeast, an enhanced correlation is observed (**Fig. 2B**). We similarly compared the folding specificity of all pairs of orthologous ncRNA genes with necessary data in yeast and mouse, including 10 snoRNA, 2 rRNA and 1 snRNA. We found strong correlation (Pearson’s *R*=0.86, *P*=0.006) despite that the sequence conservation of snoRNA is so poor that their ortholog identification has to rely on their targets (Yoshihama, et al. 2013). These results suggest that the folding specificity of a gene is moderately conserved between orthologous and paralogous gene pairs, and thus is at least partially controlled by purifying selection. Together, above observations imply that folding specificity is a gene-specific molecular trait with probable functional and evolutionary effects.

**Figure 2.**
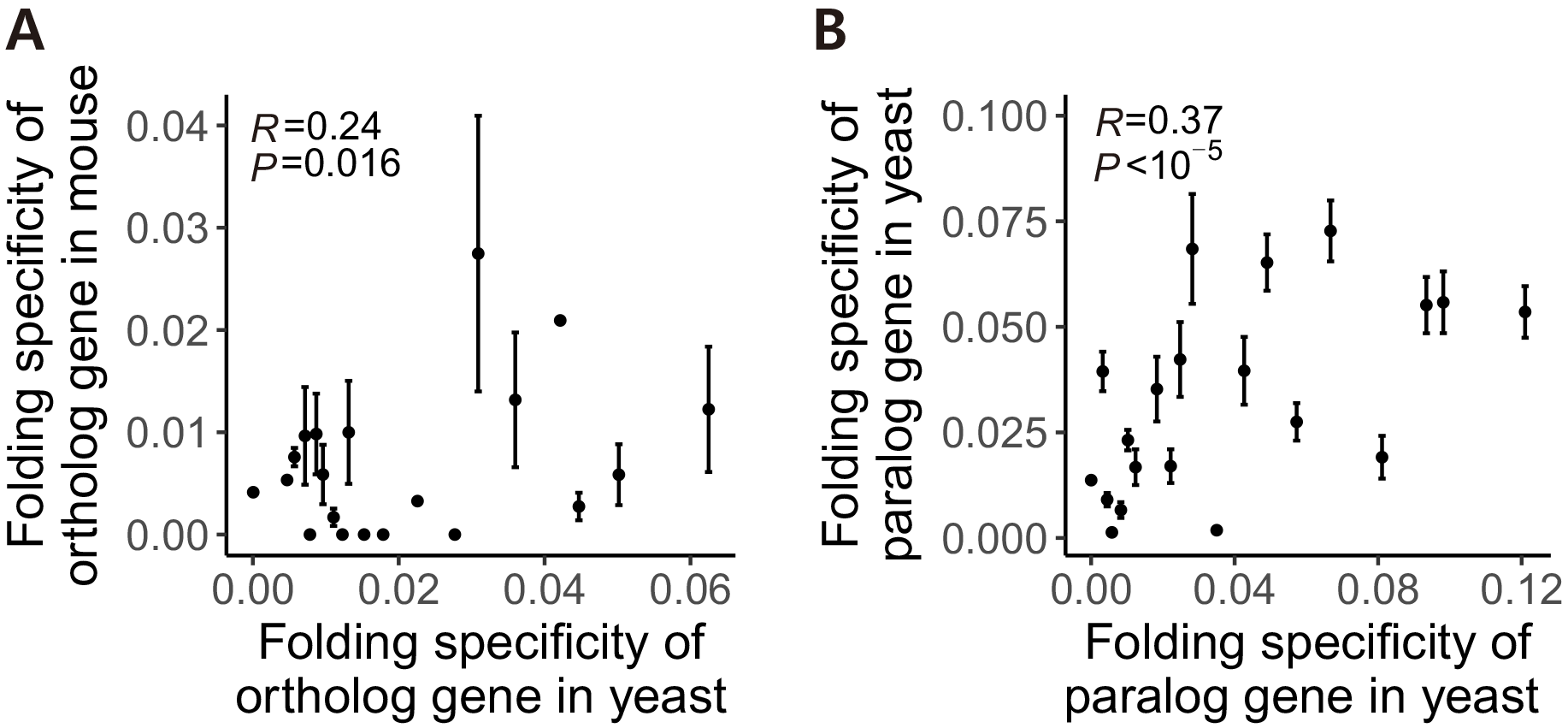
Folding specificity is an evolutionarily conserved molecular trait. (A) The folding specificity of a gene in yeast and that of the one-to-one ortholog in mouse exhibits a moderate positive correlation. All gene pairs with necessary information were divided into 20 groups by the folding specificity of the yeast ortholog (x-axis). The average folding specificity of the mouse ortholog, as well as the standard error was presented (y-axis). (B) Folding specificities of paralogous gene pairs in yeast are also positively correlated. All gene pairs with necessary information were binned into 20 groups by the folding specificity of one of the yeast paralogs (x-axis). The average folding specificity of the other paralog, as well as the standard error was presented (y-axis). Pearson’s correlation by unbinned data was also indicated within each panel.

### Relationship between folding specificity and thermostability

Thermodynamic equilibrium dictates that for any ensemble of RNA molecules with identical sequence, the fraction of molecules folded into a certain structure is exponentially proportional to the thermostability of the structure, i.e., the Boltzmann distribution. In other words, RNA has a higher probability of folding into thermodynamically more stable structure(s), which might enhance folding specificity. To find out the relationship between folding specificity and thermostability, we obtained the average melting temperatures (*T*_*m*_) of *in vitro* RNA secondary structures for each yeast mRNA, as derived from two different experimental techniques, namely DMS-seq(Rouskin, et al. 2014) and PARTE(Wan, et al. 2012) (See Materials and Methods). We divided yeast genes into two equal-sized groups with respectively high and low average *T*_*m*_. For both DMS-seq and PARTE-derived *T*_*m*_, we cannot find statistically significant difference in folding specificity between the two groups of genes (**Fig. 3A** and **B**). These results suggest that the folding specificity of mRNA is not dominated by its thermostability, which is consistent with previous observation (Zadeh, et al. 2011). Instead, it is compatible with a model where frequent unfolding by translating ribosomes, in combination with the relatively slow kinetics of exchange between alternative structures (Uhlenbeck 1995; Treiber and Williamson 1999; Woodson 2000), cause substantial deviation from the ribosome-free thermodynamic equilibrium. This phenomenon is similar to the nascent RNA emerging from RNA polymerase, which has been shown to be folding sequentially *in vivo* (Pan, et al. 1999; Heilman-Miller and Woodson 2003; Pan and Sosnick 2006; Yakhnin, et al. 2006; Wong, et al. 2007).

**Figure 3.**
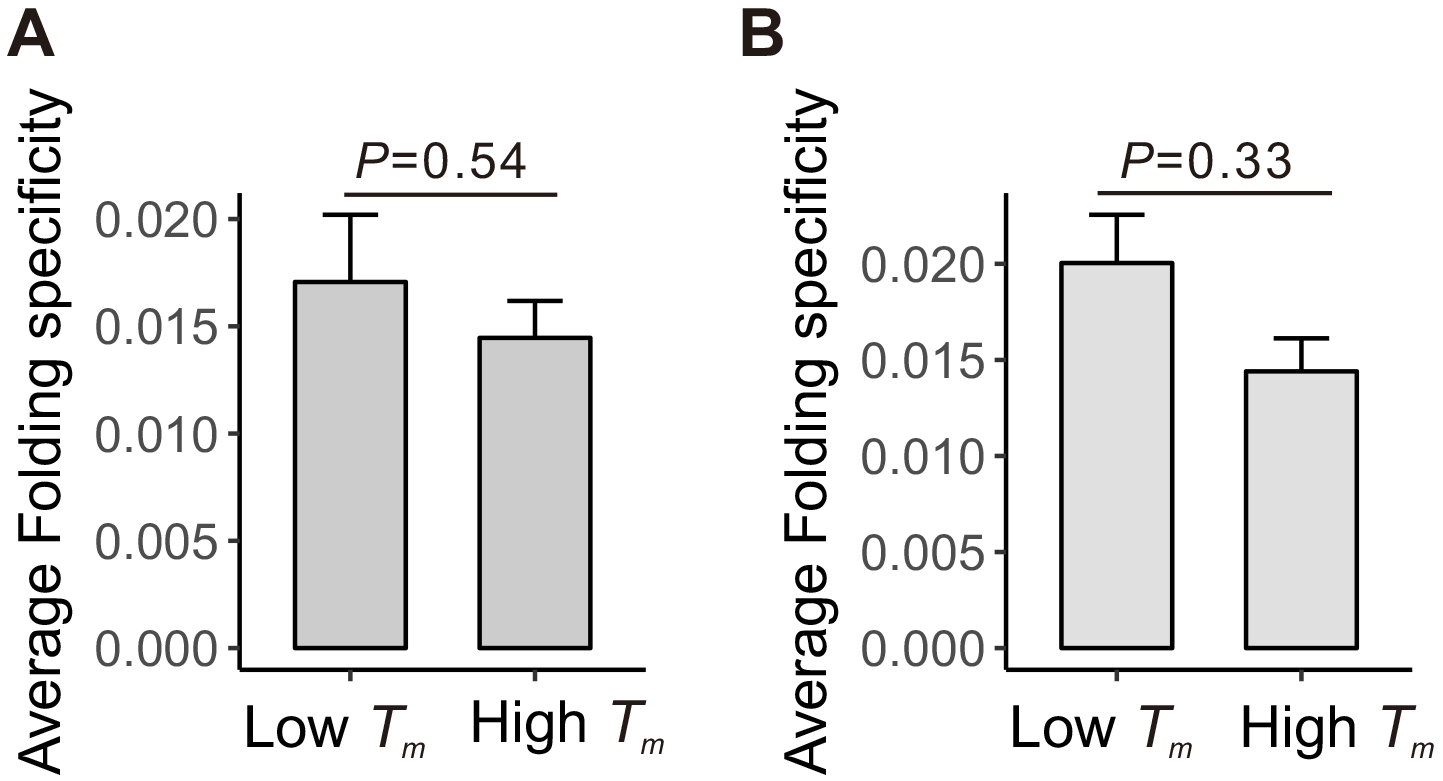
The folding specificity of mRNA is not dominated by thermodynamics. (A) Yeast protein coding genes were divided into two equal-sized groups by the melting temperature (*T*_*m*_) determined by PARTE experiment. The high *T*_*m*_ and low *T*_*m*_ group represent the 50% genes with highest and lowest *T*_*m*_, respectively. The average folding specificity is not significantly higher in mRNAs with high *T*_*m*_ *versus* those with low *T*_*m*_. (B) Same as A, except that *T*_*m*_ determined by DMS-seq is used. In both panels, error bars indicate one standard error, and *P* values are from Mann Whitney U test.

### Important genes have stronger folding specificity

Pervasive alternative folding, some evolutionarily conserved, are previously found (Lu, et al. 2016). It was argued that at least some alternative foldings are likely functional (Lu, et al. 2016). However, it remains untested whether majority of the alternative foldings, or foldings with even higher diversity, are evolutionarily adaptive. We reasoned that functionally detrimental mutations are more deleterious if happened on important genes than on other genes, therefore the adaptive molecular phenotype, be it folding specificity or diversity, should be more constrained by (purifying) natural selection in important genes. In other words, folding specificity should be more pronounced in important genes if it is generally adaptive, or vice versa for folding diversity. To test this, we compared folding specificity with different proxies of gene importance.

First, gene importance can be measured by gene indispensability, i.e., the opposite of the organismal fitness upon deletion of a gene. We estimated the indispensability of a gene by negative of the fitness of the yeast strain that lacks the gene (Steinmetz, et al. 2002), and compared it with folding specificity of corresponding mRNAs. As a result, we found that folding specificity is positively correlated with gene indispensibility (Spearman’s rank correlation ρ = 0.11, *P* = 0.009. **Fig.4A**), with two-fold higher folding specificity for the 5% most important compared to the 5% least important genes. In mouse, there is no gene indispensability data. We thus divided mouse genes into essential and non-essential groups, and found that folding specificity is significantly higher for essential than non-essential genes (*P* < 10^−7^, Wilcoxon rank sum test. **Fig. S3A**). These results suggest that folding specificity is generally adaptive, and that folding diversity is likely non-adaptive molecular or experimental noise.

Second, we used mRNA expression level as a proxy of gene importance. It is believed that due to the sheer number of the mutant molecule, mutations in highly expressed mRNA exert cytotoxicity that is otherwise negligible in lowly expressed mRNA (Zhang and Yang 2015). If folding specificity plays a role in repressing such expression-dependent cytotoxicity, natural selection should have maintained high folding specificity in highly expressed genes. Consistent with the pattern in gene indispensability, we found that folding specificity is positively correlated with mRNA expression level in yeast, where the 5% most abundant mRNAs display four-fold higher folding specificity compared to the 5% least expressed genes (Spearman’s rank correlation ρ = 0.20, *P* < 10^−6^. **Fig. 4B**). To further examine whether this pattern is an artefact created by the abundance of chimeric reads for highly expressed mRNA, we randomly sampled five intramolecular chimeric reads from each mRNA and recalculated folding specificity. Such randomized down-sampling was repeated for 1,0 times, and the resulting folding specificity was still positively correlated with mRNA expression level (*P* = 0.039, permutation test. **Fig. S3D**). In addition, we compared the folding specificity with mRNA expression level in mouse, and again identified positive correlations (Spearman’s rank correlation ρ = 0.28, *P* < 10^−127^. **Fig. S3B** and **E**), lending further support to the adaptiveness of folding specificity.

**Figure 4.**
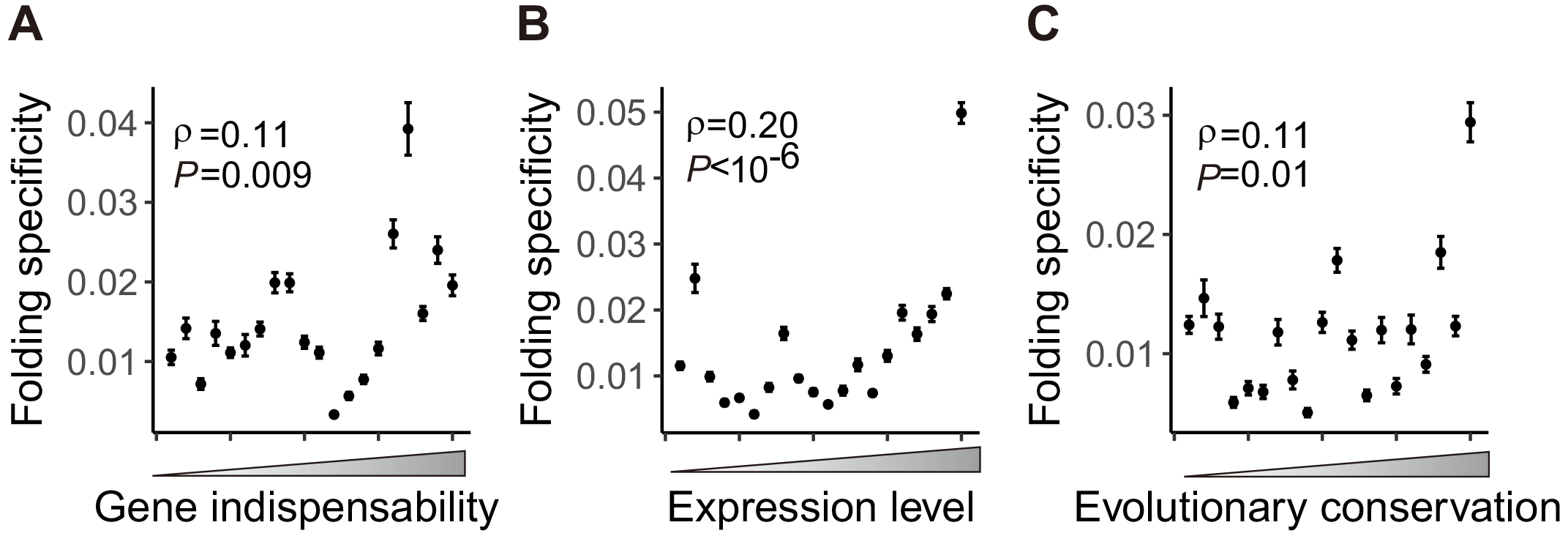
Important genes have stronger folding specificity. The functional importance of a gene is approximated by gene indispensability (A), mRNA expression level (B) and evolutionary conservation (C). The genes were binned into 20 equal-sized groups by their rank in the three proxies of functional importance. The mean folding specificity of each group, as well as its 95% confidence interval assessed by 1,000 bootstrapping of the genes, were indicated by the dots and the corresponding error bars. The Spearman’s correlation coefficients of the unbinned data are also shown. All three panels consistently shown that, the folding specificity of a gene increases with its functional importance.

Third, we compared folding specificity with the evolutionary conservation (See Methods) of the gene. Because highly conserved genes are under stronger functional constraint (Koonin and Wolf 2010; Zhang and Yang 2015), we shall predict stronger folding specificity for more conserved than less conserved genes, given previous observation made by protein indispensability and expression level. Indeed, we found a positive correlation between evolutionary conservation and folding specificity in both yeast (Spearman’s rank correlation ρ = 0.11, *P* = 0.01. **Fig. 4C**) and mouse (Spearman’s rank correlation ρ = 0.14, *P* < 10^−29^. **Fig. S3C**). In summary, three different proxies of gene importance consistently support that folding specificity is adaptive, and on the other hand suggest that folding diversity is likely nonadaptive phenomenon derived from molecular stochasticity or experimental noise.

### Increased folding specificity of highly expressed mRNAs is not caused by RNA circulation due to selection for translational efficiency

Hereinafter, we will focus on RPL data from yeast unless otherwise noted, because of the availability of various types of functional genomic data in yeast (see below), and its relatively higher coverage for more accurate quantification of folding specificity. It was previously reported that circularization of mRNA by eukaryotic translation initiation factors facilitates ribosomal recycling and efficient mRNA translation (Wells, et al. 1998). Indeed, RNA folding with long intervening distance is more prevalent in genes with high translation efficiency than those with low translational efficiency (Aw, et al. 2016). If highly expressed mRNAs are also highly translated (Muzzey, et al. 2014), the observed correlation between mRNA expression level and folding specificity might then be explained by the dominance of long distance folding, in particular those connecting 5’ and 3’ ends of the mRNA. To rule out such possibility, we calculated the “circularization score” (Aw, et al. 2016) for each RNA folding, which is the distance between the center nucleotides of the two folding partners supported by each chimeric read, normalized by the gene length. We then used 5% of chimeric reads with top (distal folding) or bottom (proximal folding) circularization score to recalculate folding specificity. The correlation between folding specificity and mRNA expression level is significantly positive when we use this subset of the data, and remains so if we included up to 50% of distal/proximal folding chimeric reads (**Fig. 5**). The above result indicates that RNA foldings of various distances all contribute to the higher folding specificity of highly expressed mRNAs, which is thus unexplainable by the RNA circulation due to selection for translational efficiency. Furthermore, our observation suggests that, instead of a local feature limited to certain fraction of the mRNA sequence (such as 3’ end, 5’ end, UTR, etc), folding specificity impacts the whole mRNA molecule.

**Figure 5.**
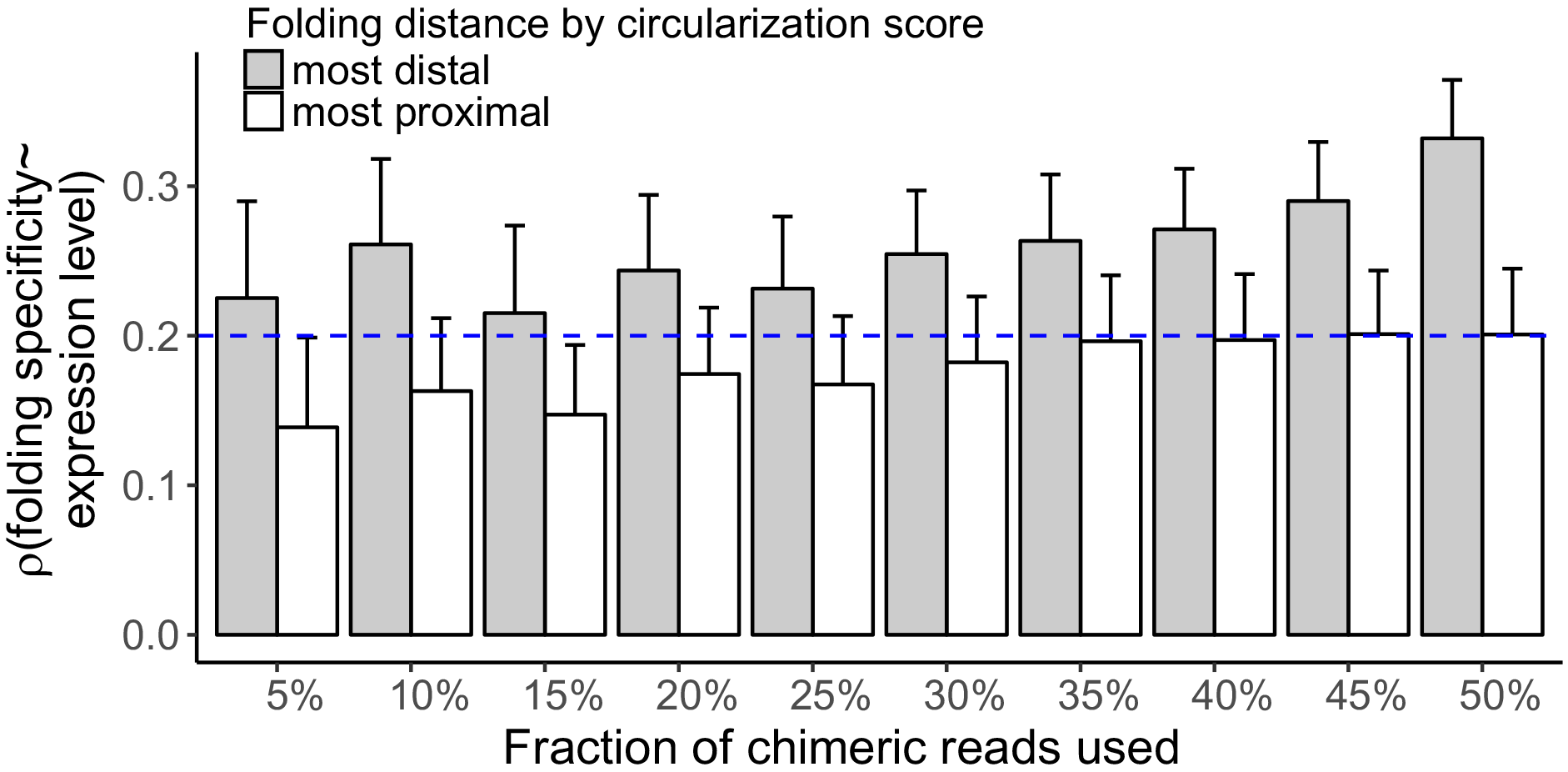
Increased folding specificity of highly expressed mRNAs is not caused by RNA circulation due to selection for translational efficiency. Chimeric reads and their indicated foldings were categorized into proximal/distal folding by their circulation score. The spearman’s correlations between mRNA expression level and the folding specificity estimated by using different fractions of proximal or distal foldings in every gene are shown. Error bars indicate one standard error, estimated by bootstrapping the genes 1,000 times. The blue dashed line indicates the correlation when all intramolecular foldings were used

### Conserved nucleotides fold more specifically than less conserved nucleotides within the same gene

With above results showing positive correlation between folding specificity and gene importance, prediction can be similarly made that within the same gene, specific foldings should be associated with important region of the gene. Furthermore, within-gene comparison of functional importance with folding specificity is completely free of intergenic confounding factors such as expression level. To this end, we calculated folding specificity for each nucleotide of an mRNA using the chimeric reads supporting folding for the focal nucleotide (See Materials and Methods. **Fig. 6A**).

**Figure 6.**
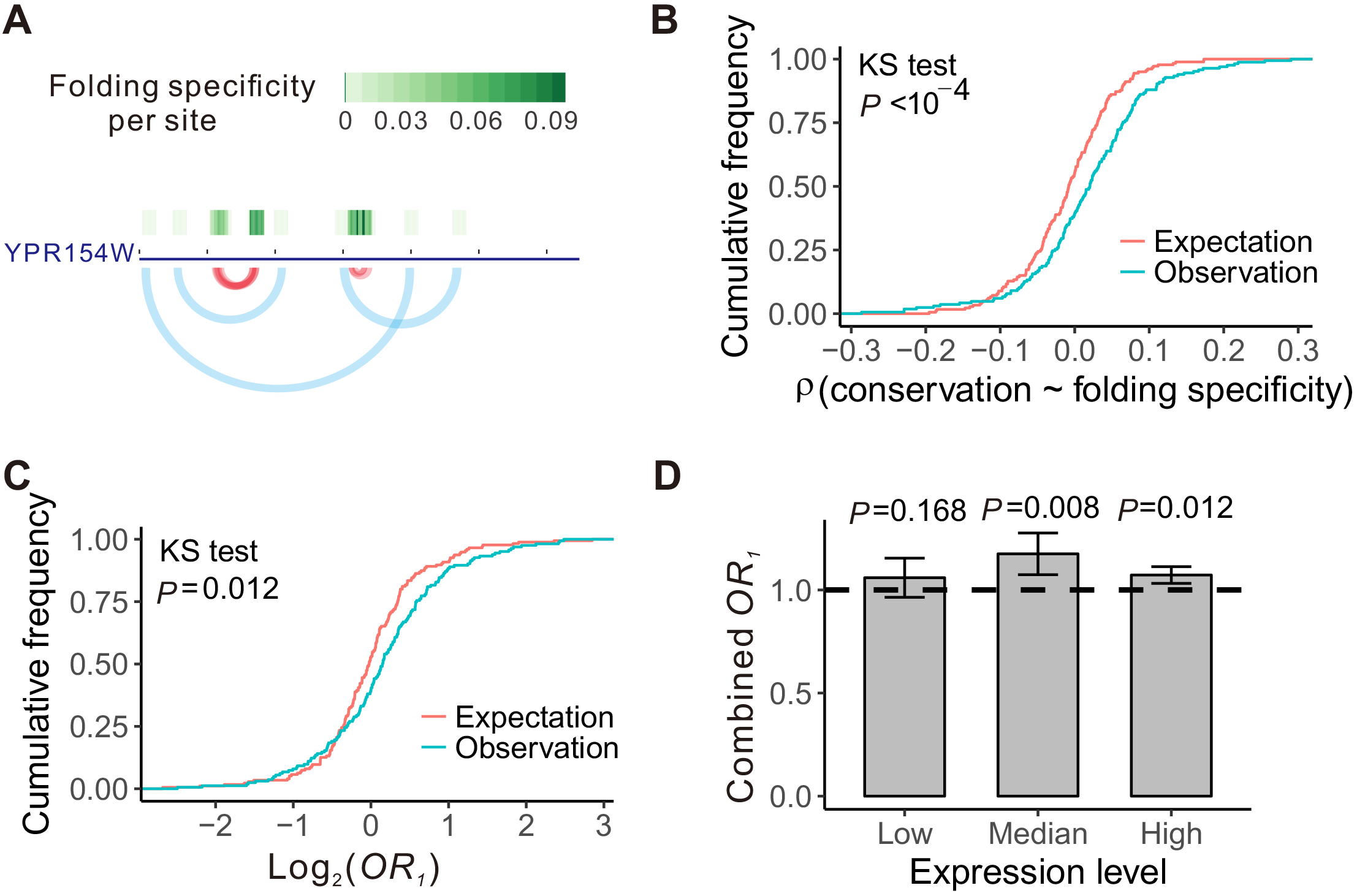
Stronger folding specificity for more conserved nucleotides within gene. (A) An example showing folding specificity calculated for each nucleotide. The gene and the arches indicating foldings are same as Fig.1A. The per-site folding specificities were further shown as green tiles above the gene, whose levels are indicated by the color scale. (B) Cumulative frequency distribution for the observed and randomly expected within-gene spearman’s correlations between the evolutionary conservation of each nucleotide and its folding specificity. The random expectation is generated by shuffling per-nucleotide conservation level. *P*-values are from Kolmogorov-Smirnov(KS) test. (C) Cumulative frequency distributions for the observed and randomly expected odds ratio (*OR*_1_), which measures the enrichment of nucleotides with higher folding specificity at evolutionary conserved residues within a gene. The random expectation is generated by shuffling per-nucleotide conservation level. *P*-values are from Kolmogorov-Smirnov test. (D) The yeast genes analyzed were grouped into three equal-sized bins according to their expression level, and their combined *OR*_1_ values estimated by Mantel-Haenszel procedure are shown. All three groups display *OR*_1_ > 1 (the horizontal dashed line), indicates that conserved nucleotides have higher folding specificity than less conserved ones within a gene, although the trend is not statistically significant in lowly expressed genes. Error bar indicates one standard error, estimated by bootstrapping the genes 1,000 times.

We then reasoned that the functional importance of each nucleotide can be approximated by its evolutionary conservation, which was estimated using one-to-one orthologs in 6 post-WGD (whole genome duplication) yeast species (See Materials and Methods). The level of evolutionary conservation and folding specificity for each nucleotide was subsequently compared for each of 166 distinct genes with necessary information. Consistent with our prediction, positive Spearman’s rank correlation was found in 101 genes, significantly more than the random expectation of 166/2=83 (*P*= 0.006, binomial test). For each gene, we also randomly shuffled the folding specificity of each site and re-evaluated the correlation between evolutionary conservation and folding specificity, which serves as an *ad hoc* random expectation. Compared with this expected distribution, the real within-gene correlation between folding specificity and conservation is significantly skewed towards positive values (**Fig. 6B**), suggesting association between important nucleotide and specific folding, and that specific folding is likely more adaptive than non-specific folding.

To further assess the relationship between evolutionary conservation and folding specificity for each gene, we constructed a 2 × 2 matrix for each gene by respectively dividing each site of the gene into one of four groups on the basis of its folding specificity and evolutionary conservation, and calculated an odds ratio (*OR*_1_) from the matrix (see Materials and Methods). If the specifically folded sites are preferentially located at conserved regions, the *OR*_1_ is > 1. We similarly generated a randomly expected *OR*_1_ distribution by shuffling the folding specificity among all sites within each gene, which was found dwarfed by the real *OR*_1_ values (**Fig. 6C**). This result again supports the adaptiveness of specific folding.

To determine whether the conservation level at specifically folded sites was affected by expression level, we divided genes with necessary information into three groups with the low, median and high expression level, and calculate an overall *OR*_1_ for each group using the Mantel-Haenszel (MH) procedure (See Materials and Methods and **Fig. S4**). As expected, in all cases except the low expression level, the combined *OR*_1_’s significantly exceed 1 (**Fig. 6D**), suggesting that conserved sites in highly expressed genes display stronger propensity to fold specifically, consistent with the stronger selection for highly expressed genes. Finally, we combined all genes for an overall *OR*_1_ = 1.09 (*P*<10^−4^, MH test), again lends support for the adaptiveness of folding specificity. This observed association between conservation and folding specificity within gene appears unexplainable by the local thermodynamic stability, because nucleotide-wise *T*_*m*_ has no effect on the per nucleotide folding specificity, as evident by the insignificant *OR*_2_ (See Materials and Methods) testing overrepresentation of nucleotides with low *T*_*m*_ and high folding specificity regardless whether DMS-seq or PARTE-derived *T*_*m*_ is used (**Fig. S5**). In combination with the observation made by between gene analyses, our results offered unequivocal support for an overall adaptive role for folding specificity, and suggested a non-adaptive role, and thus likely molecular or experimental noise, for folding diversity.

### The Case of Ribosome Stalling Demonstrates Functional Impact of Folding Specificity

Given the results presented above, we then ask the question: what is the molecular mechanism that grants folding specificity its selective advantage? As we’ve shown the adaptiveness of folding specificity regardless the folding distance (**Fig. 5**), the functional benefit conferred by specific folding is likely generally applicable to the whole mRNA molecule, instead of confined to a small specific region such as 5’ and 3’ ends (e.g. regulating initiation rate(Kudla, et al. 2009)) or intron/exon borders (e.g. regulating alternative splicing(Graveley 2005)). Therefore, we chose to test the functional impact of folding specificity on ribosome stalling, a molecular phenomenon that is potentially applicable to the whole body of CDS (Tuller, et al. 2011; Yang, et al. 2014). Nevertheless, it is by no means indicating that this is the only functional benefit provided by specific folding.

We have previously shown that ribosome stalling by mRNA secondary structure modulates translational elongation speed, which is likely utilized by natural selection to balance the trade-offs between translational accuracy and efficiency (Yang, et al. 2014). Other reports also suggested a regulatory role for mRNA secondary structure in co-translational protein folding (Yang and Zhang 2015; Faure, et al. 2016; Yang 2017). We thus asked whether folding specificity affects the capacity of RNA secondary structure in stalling upstream ribosomes. To this end, we compared the first (closest to 5’ end of the mRNA) nucleotide showing highest folding specificity within each gene, with expression-normalized local Ribo-Seq coverage (See Materials and Methods). We averaged, across genes, the normalized ribosome densities near the specifically folding nucleotides, and found a significant increase of ribosomal density upstream of the site with specific folding. In particular, we found a maximum of 32% increase of ribosome density at the 42^th^ nucleotide upstream of the most specifically folded nucleotide (**Fig. 7A**, red line). This magnitude of ribosome stalling by mRNA folding is comparable to previous reports (Charneski and Hurst 2013; Yang, et al. 2014), and the position of the peak of ribosome density likely reflects the limited resolution of RPL (Ramani, et al. 2015) and Ribo-seq (Diament and Tuller 2016), but cannot be explained by the 5’ ramp of translation elongation speed (Tuller, et al. 2010), because all specifically folded nucleotides are at least 200 nucleotides downstream from translational start site.

**Figure 7.**
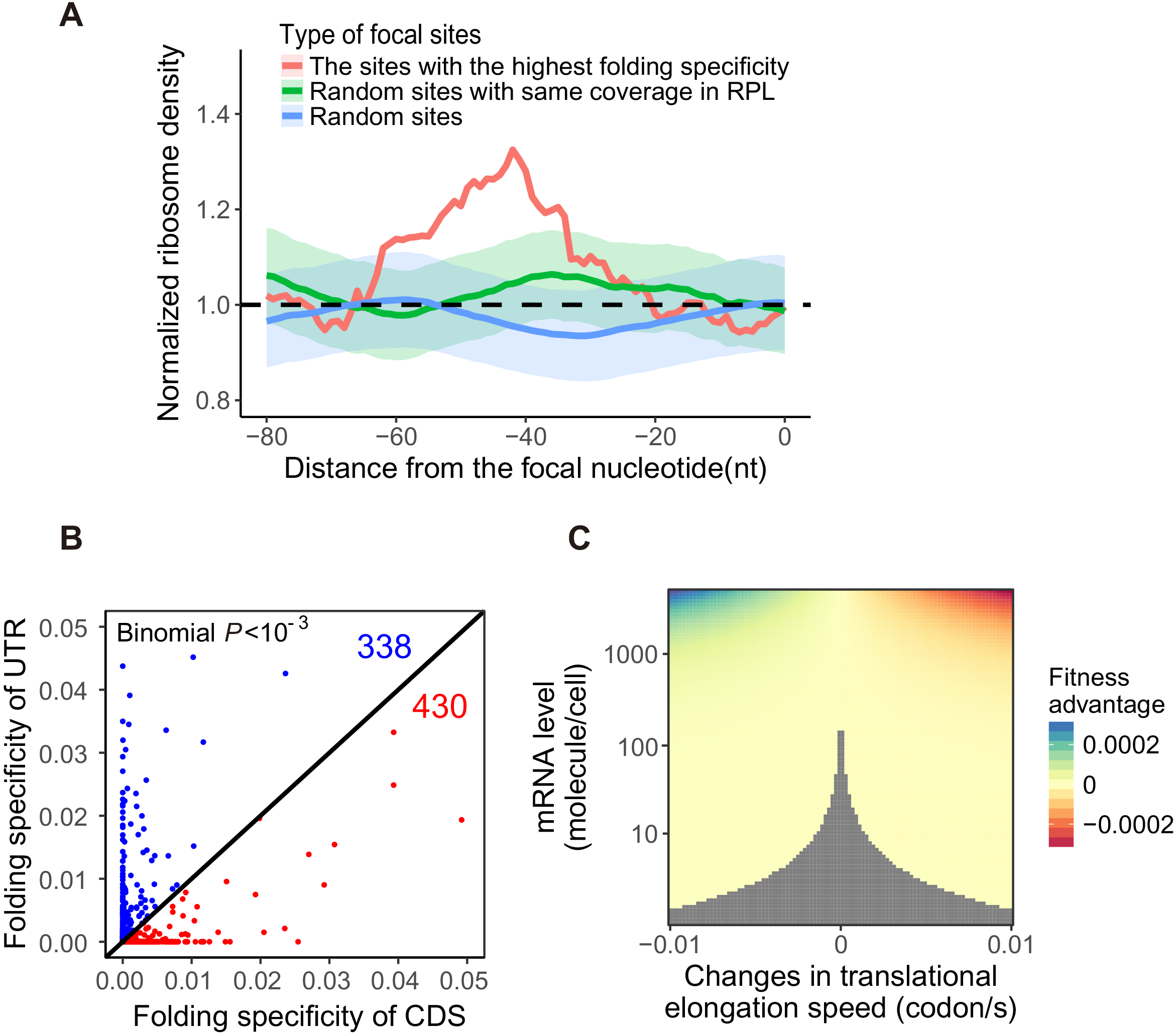
Effect of folding specificity on ribosome stalling. (A) Meta-gene analysis of normalized ribosome density for the upstream of the first most specifically folded nucleotide with *S* > 0.05 within each gene analyzed. The most specifically folded nucleotide was at *x* = 0, upstream of which a peak of ribosome density was shown (red line). As a control, another random site with non-zero coverage in RPL and *S* = 0 was chosen for each gene, and the average normalized ribosome density for the upstream of this focal site, as well as the standard error, were shown as blue line and shade. Another control, derived from random site with same RPL coverage as the most specifically folded site and *S* = 0, was similarly shown as green line and shade. Neither control display significant deviation from the dashed black line of *y* = 0, which indicates no increase of ribosome density. (B) For each gene, the per-nucleotide folding specificity of CDS is averaged (x axis) and compared with that of UTR (y-axis). The dot representing a gene will thus lie below the diagonal line if the CDS is on average more specifically folded than the UTR (red dots), or *vice versa* (blue dots). There are significantly more red dots (430) than blue dots (338) than random expectation (Binomial *P* < 10^−3^). (C) Given the influence of folding specificity on ribosome density shown in (A), the fitness effect of mutations that change folding specificity can be estimated by a model that considers ribosome sequestration and translational fidelity (See Materials and Methods). If a change of folding specificity can alter the ribosome density of one codon by ~20%, as shown in (A), the average elongation speed for a gene will changed by 0.01 codon/s, which is used as the limits of x-axis (See Materials and Methods). The fitness advantage, as indicated by the colors, is shown as a function of elongation speed (relative to the baseline of 20 codons per second) and gene expression level of the focal gene. The gray area represents fitness effects that are too small to be targeted by natural selection in yeast, whose effective population size ~ 10^7^.

To assess the expected ribosome density around a random nucleotide captured by RPL, we repeated the local ribosome density analysis using sites with nonspecific folding (i.e. *S* = 0) and found no significant increase of ribosome density upstream of the focal nonspecific site (**Fig. 7A**, blue line and shade). To further elaborate the effect of specific folding *versus* non-specific folding, we again repeated the local ribosome density analysis with sites having same number of chimeric reads as the specifically folded sites and *S* = 0 (**Fig. 7A**, green line and shade). This additional control again showed no significant increase of ribosome density upstream of non-specifically folded nucleotide, suggesting exclusive association of specificity folding with upstream ribosome stalling.

Other than ribosome stalling by specific mRNA folding, an alternative explanation for the aforementioned observation is that ribosomes limit the folding choices of flanking nucleotides by steric hindrance, effectively increasing the folding specificity. However, this alternative explanation would predict peaks for Ribo-Seq reads at both upstream and downstream of the specifically folding sites, as well as no skewed folding specificity due to gene importance. Since both predictions contradict with observation made in empirical data (**Fig. S6** and **Fig. 4**), this result supports a role of folding specificity in the ribosome-stalling capacity of mRNA secondary structure. More importantly, this result suggests that highly expressed genes, which have increased folding specificity, have better control over ribosome velocity. Such feature of highly expressed genes is consistent with its increased requirement for translational fidelity (Yang, et al. 2014) and/or co-translational protein folding accuracy (Yang, et al. 2010).

To further validate the functional role of folding specificity, we calculated for each gene the average per nucleotide folding specificity of CDS, and that of UTR. The mouse PARIS data was used here because UTR annotation is rare for yeast. If folding specificity is involved in the regulation of translation by its ribosome stalling capacity, we shall predict that the average folding specificity of CDS should be higher than that of UTR, because translation only happens on CDS. In support of our prediction, we found that for majority of mouse mRNAs, the folding specificity of CDS is stronger than that of UTR (Binomial *P* < 10^−3^. **Fig. 7B**), reinforcing the support for our hypothesized connection between folding specificity and translation.

Finally, we asked whether the functional benefit provided by specific folding and its ability to slow down elongation is strong enough to become a target of natural selection. To this end, we used a previously published model (Yang, et al. 2014), which incorporates experimentally determined parameters in yeast to predict the fitness effect of changes in translational elongation speed. This model considers two competing selective pressures, one for increased elongation speed, which reduces ribosome sequestration, and the other for reduced elongation speed, such that translational accuracy is higher (Johansson, et al. 2012; Yang, et al. 2014) and protein misfolding (O’Brien, et al. 2012) is less likely (See Materials and Methods). We found that, a mutation that induced specific folding and increase ribosome density on one codon by 20% will makes the cell 0.03% fitter if the mutation happen on highly expressed (5000 molecules/cell) genes (**Fig. 7C**, top). Because 3×10^−4^ greatly exceeds the inverse of the effective population size (10^7^) of yeast (Wagner 2005), such a mutation can be targeted by natural selection. Moreover, we found that for a lowly expressed genes (1 molecules/cell), the same mutation only corresponds to a fitness effect of *s* = 7.0×10^−8^ (**Fig. 7C**, bottom), making it effectively a neutral mutation (*s* < 10^7^). In other words, the selection for folding specificity on important or evolutionarily conserved nucleotides shall be significant in highly expressed genes, but not in lowly expressed genes, which is exactly what we observed (**Fig. 6D**). Interestingly, this result also suggests that there shall be no selection for specific folding in human, because even happened in highly expressed genes, the selective coefficient for enhanced folding specificity (3×10^−4^) is still too small compared to the inverse of the human effective population size (~10^3^) (Park 2011), which is exactly what we observed using folding specificity derived from human PARIS data (Lu, et al. 2016) (data not shown).

Altogether, this result suggest that folding specificity of a mRNA can be potential targets of natural selection depending on the mRNA expression, which corroborates the above observation of weaker folding specificity for lowly expressed genes compared to highly expressed genes.

## Discussion

In this study, we utilized recently published high throughput sequencing data for RNA duplexes to estimate the specificity of mRNA folding. Consistent with the disruption by translating ribosome, we found the folding specificity of mRNA to be significantly lower than that of noncoding RNA. Unexpectedly, the folding specificity is not stronger for secondary structures with higher thermostability *in vitro*. We further found a positive correlation between folding specificity and functional importance among genes and among sites within the same gene, suggesting an evolutionarily adaptive role of specific folding. In search of the molecular function underlying the benefit of specific folding, we contrasted nucleotides with specific *versus* promiscuous folding, and revealed the capacity of ribosome stalling for specific but not promiscuous folding. Collectively, our results demonstrated the evolutionary and functional significance of folding specificity, and offered new insights for the study of mRNA secondary structure.

One potential caveat in our analyses is that the estimation of folding specificity might be confounded by the number of short reads supporting any folding partnerships within each gene. By incorporating multiple methods, including comparison with theoretically maximal entropy, down-sampling chimeric reads to a unified number, and within-gene comparison, we showed that the correlation between functional importance and folding specificity is robust regardless such confounding factor. On the contrary, the limited resolution of, as well as the technical/organismal difference between yeast RPL and mouse PARIS data potentially add random noise to the actual biological signal, which is thus likely stronger than shown in our study.

The level of RNA folding specificity is expected to be influenced by RNA binding proteins. For non-coding RNAs, proteins likely stabilize the native functional RNA foldings, and thus increase folding stability. On the contrary, mRNA foldings are expected to be constantly disrupted by translating ribosomes. Indeed, RNA folding occurs on a microsecond time scale(Gralla and Crothers 1973; Porschke 1974), which is much faster than the <30 codons per second translational elongation rate in vivo(Gilchrist and Wagner 2006; Ingolia, et al. 2011). Therefore, mRNA regions that are not occupied by ribosomes have enough time to form local secondary structures, which should change as ribosomes move. Maintaining specific folding in the face of such frequent disruptions suggests tight regulation for the structure, which is therefore likely functional.

Assuming thermodynamic equilibrium, RNA folds into various secondary structures with probabilities dictated by the folding energies. It is therefore possible that folding specificity simply reflects the thermostability of the RNA molecule. In opposite to this possibility, we found no correlation between folding specificity and average melting temperature(Wan, et al. 2012) of RNA secondary structure for a gene. As additional support to our finding, previous study on RNA design has already shown that stability and specificity are poorly related (Zadeh, et al. 2011). The discrepancy between folding specificity and thermostability may be explained by the effect of translation on mRNA, where ribosomal occupation allows sequential local foldings but excludes thermodynamically-favored global foldings. This phenomenon is similar to the nascent RNA emerging from RNA polymerase, which has been shown to be folding sequentially in vivo (Pan, et al. 1999; Heilman-Miller and Woodson 2003; Pan and Sosnick 2006; Yakhnin, et al. 2006; Wong, et al. 2007).

The diversity of RNA folding, as inversely approximated by folding specificity, appears evolutionarily non-adaptive according to comparison with functional importance of genes or sites within the same gene. This is compatible with a model where molecular stochasticity, a largely non-adaptive intrinsic property underlying all biological processes, has been selectively reduced for important genes. This model is supported by multiple biological phenomenon at molecular level, such as protein expression noise (Metzger, et al. 2015), misinteraction (Yang, et al. 2012), misfolding (Yang, et al. 2010), etc(Park and Zhang 2011; Xu and Zhang 2014; Liu and Zhang 2017a, b). According to this model, alternative RNA secondary structure, especially for mRNA, is likely non-adaptive, and selectively constrained by purifying selection against molecular stochasticity. Note, however, that a small fraction of alternative folding, especially those with relatively high folding specificity, might still be conserved and functional (Lu, et al. 2016).

We have found intergenic and intragenic evidence for the evolutionary adaptiveness for folding specificity. There are several hypotheses regarding the exact functional benefit provided by specific mRNA folding that are worthy of discussion here. First, Mao and colleagues(Mao, et al. 2014), based on computational simulation, proposed that strong mRNA folding without ribosomes slows down the first translating ribosomes, thereby shortens distance between subsequent ribosomes and eliminates secondary structure in translating mRNA in increased ribosomal occupancy, effectively increasing the translational efficiency. Under this model, specific foldings might have played important role in slowing down the first translating ribosome. However, we found this model by Mao and colleagues paradoxical because slowing down the first translating ribosome is expected to decrease instead of increase translational efficiency. Moreover, it has been estimated that ~70% of coding regions are unoccupied by ribosomes (Arava, et al. 2003; Zenklusen, et al. 2008), which is more than enough available nucleotides for translating mRNA to fold, given that the rate of RNA folding (in microseconds(Gralla and Crothers 1973; Porschke 1974)) is much faster than ribosomal translocation (< 30 codons per second(Gilchrist and Wagner 2006; Ingolia, et al. 2011)). Indeed, the RNA duplexes found by RPL(Ramani, et al. 2015) and PARIS(Lu, et al. 2016) is consistent with rich secondary structure in translating mRNA. Therefore this model is an unlikely explanation for the functional benefit of specific folding.

Another hypothesis by Qi and Frishman (Qi and Frishman 2017) proposed that RNA secondary structure with high and low thermostability are under evolutionary pressure to preserve RNA secondary structure and primary sequence, respectively. This model might partially explain the functional benefit provided by specific folding if folding specificity is correlated with thermostability. However, we found genes with stronger folding specificity are not thermodynamically more stable. Therefore this model cannot provide a mechanistic link between folding specificity and its evolutionary adaptiveness.

A third hypothesis is that specific RNA folding serves as a molecular brake on translating ribosomes, thus enhancing the fidelity of translation and/or co-translational protein folding(O’Brien, et al. 2012; Shalgi, et al. 2013; O’Brien, et al. 2014; Yang, et al. 2014). Indeed, it was suggested that mRNA structure acts as a gauge of co-translational protein folding by reducing ribosome speed when extra time is needed by the nascent peptide to form and optimize the core structure(Faure, et al. 2016). This model is compatible with a considerable body of experimental evidence indicating synonymous variants capable of (de-)stabilizing mRNA secondary structures can dramatically alter translation speed and influence co-translational protein folding(Nackley, et al. 2006; Kimchi-Sarfaty, et al. 2007; Komar 2007; Shabalina, et al. 2013). According to our result, specific folding is more efficient in regulating ribosome speed.

The mechanism behind this effect of folding specificity remains to be answered, because specific and non-specific foldings should be similar obstacles for ribosome movement, especially when specific folding is not thermodynamically more stable. One potential explanation is that specific folding is more resistant to the helicase activity of ribosome (Takyar, et al. 2005) than non-specific folding *in vivo*, such that resolving specific folding requires extra time. Nevertheless, regardless the molecular mechanism, the increased folding specificity in highly expressed genes is consistent with their stronger tendency to avoid mistranslation (Drummond and Wilke 2008; Yang, et al. 2014) and misfolding (Yang, et al. 2010), which imposes an expression-dependent fitness cost (Drummond and Wilke 2008; Geiler-Samerotte, et al. 2011).

While significant advances in high throughput experimental technique have enabled dissection of RNA secondary structures on transcriptome scale, extracting functionally relevant RNA foldings have remained challenging, especially for mRNA, which are constantly disrupted by translating ribosomes. The positive correlation between folding specificity and functional importance among genes and sites within the same gene, as shown in the current study, points to a novel strategy of prioritizing mRNA structures by their folding specificity. Indeed, we showed ribosome stalling at upstream of nucleotides with specific folding, but not those with promiscuous folding, demonstrating the usefulness of folding specificity on distinguishing functional vs non-functional RNA structures.

In summary, the conception of folding specificity of RNA secondary structure and its application reveal previously unappreciated complexities underlying RNA secondary structure *in vivo*. Specific folding of mRNA despite of frequent disruption by translating ribosomes appears selectively maintained, and associated with evolutionarily adaptive molecular function such as regulation of co-translational protein folding. Accounting for folding specificity shall provide valuable information in the functional study of RNA secondary structures, particularly for mRNA.

## Materials and Methods

### Genome, annotation and comparative genomic data

Genome and transcript sequences and annotation were obtained from EnsEMBL release 89 (Aken, et al. 2017), the specific genome versions are R64-1-1 for *S. cerevisiae* and GRCm38 for *M.musculus*. The list of one-to-one orthologs between the two species were also downloaded from EnsEMBL. Each gene was represented by its longest annotated transcript. To estimate the evolutionary rate of *S. cerevisiae* mRNA, we also collected the mRNA sequences from five other post-WGD (whole-genome duplication) fungal species (*S. paradoxus*, *S. mikatae*, *S. bayanus*, *Candida glabrata*, and *S. castellii*), along with gene orthology/paralogy information among the six species from the Fungal Orthogroups Repository (Wapinski, et al. 2007). The orthologs between yeast and mouse snoRNA were extracted from snOPY database(Yoshihama, et al. 2013), where the target genes of the snoRNA were used to identify orthologs.

### High throughput sequencing data of RNA folding and the estimation of folding specificity

We used datasets derived from two distinct experimental techniques to assess RNA folding. On the one hand, the single nucleotide folding anchors for each folding partners derived from RNA proximity ligation (RPL) (Ramani, et al. 2015) assay in yeast *S.cerevisiae* were downloaded from the NCBI Gene Expression Omnibus (GEO) (Barrett, et al. 2013) under accession number GSE69472. Only the intramolecular folding pairs were retained for further analysis. On the other hand, the raw reads from psoralen analysis of RNA interactions and structures (PARIS) (Lu, et al. 2016) conducted in mouse were downloaded under accession number GSE74353. The raw reads were then processed using analytical pipelines provided by the authors (https://github.com/qczhang/paris) (Lu, et al. 2016) to yield a list of folding partners. Briefly, short reads were adaptor-trimmed and merged, and then aligned to the genome (mm10) by STAR aligner(Dobin, et al. 2013). The reads mapped with gap or chiastically were combined and assembled into duplex groups by a two-step greedy algorithm, as implemented by the scripts provided by the authors(Lu, et al. 2016). Finally, short reads were extracted from intramolecular duplex groups, and the nucleotides in the center of 5’ or 3’ fragment mapped to either folding partner are used as anchors for folding partnership.

Due to the limited resolution, it is difficult to locate the exact pairing partner from RPL or PARIS data. Following the bioinformatics analyses in RPL assay(Ramani, et al. 2015), we instead generated contact probability maps using anchors of folding partnership derived from either RPL or PARIS. Briefly, we used python scripts by RPL authors (Ramani, et al. 2015) and computed the coverage at each base *i* and *j* (*C*_*i*_; *C*_*j*_) and generated a normalized matrix *M*_*norm*_ such that 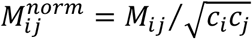, where *M*_*i,j*_ is the number of reads supporting folding partnership between *i* and *j*. We then used this matrix to generate *M*\* by binning normalized scores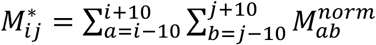. Effectively, this is indicating support for folding partnership for the 21 nucleotides (anchor ± 10 nts) linearly surrounding any pair of folding anchor.

On the basis of 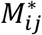, we can calculate the folding specificity for a whole gene. Adapted from Shannon entropy(Huynen, et al. 1997), we first calculate *S*_*obs*_ = −Σ_*i,j*_*p*_*i,j*_ln*p*_*i,j*_ for a gene *g*. Here, *p*_*i,j*_ is the relative folding probability between nucleotide *i* and *j*, or in other words 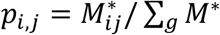, with Σ_*g*_*M*\* being the sum of all 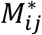 within *g*. To guard against the confounding effect of sequencing depth, we also calculated the theoretical maximal of *S*_*obs*_ as 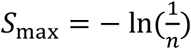, where *n* is the total number of pairs of nucleotides whose physical proximity are revealed by at least one chimeric read. Here *S*_max_ equals to the information entropy when folding of every relevant bases are equally supported. We then calculate folding specificity as *S* = *S*_max_ − *S*_obs_, where higher *S* represents stronger folding specificity. The formula for *S* is mathematically equivalent to Theil index, a commonly used metric for economic inequality(Cowell 2003).

Similarly, we can calculate *S* for a (not necessarily continuous) region of a gene by defining 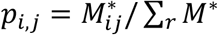, with Σ_*r*_*M*\* being the sum of all 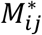 within *r*. We also calculated the folding specificity for individual nucleotides by defining 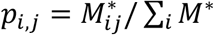, where *i* is the focal nucleotide under study and Σ_*i*_*M*\* is the sum of all 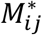 involving *i*.

### Thermostability of mRNA secondary structure

We estimated the thermostability of yeast mRNA secondary structure using data from two experimental techniques, namely PARTE (Parallel Analysis of RNA structures with Temperature Elevation)(Wan, et al. 2012) and DMS(dimethyl sulphate)-seq (Rouskin, et al. 2014). In PARTE, footprinting of double-stranded RNA residues by RNase V1 across five temperatures (from 23 to 75 °C) was coupled with high-throughput sequencing to reveal the energetic landscape of the transcriptome(Wan, et al. 2012). In DMS-seq experiments, modification of unpaired adenine and cytosine by DMS was monitored by deep sequencing. DMS-seq was used to evaluate the *in vitro* thermostability of RNA folding by genome-wide DMS-seq assays in five temperatures (from 30 to 95°C) (Rouskin, et al. 2014). In both PARTE and DMS-seq, RNA secondary structure unfolds as temperature rises, giving rise to estimation for the melting temperature (*T*_*m*_).

We downloaded PARTE and DMS-seq data from GEO under accession number GSE39680 and GSE45803, respectively. The raw reads for both datasets were adaptor-trimmed and mapped to yeast genome, followed by *T*_*m*_ estimation using previously published computational procedure (Wan, et al. 2012). Briefly, the data were normalized by the library sizes estimated by PossionSeq (Li, et al. 2012) and then fitted to an adaptive regression model to search for sharp transitions in read numbers at each probed base as a function of temperature(Wan, et al. 2012). We then averaged *T*_*m*_ for all nucleotides with necessary information to represent the average thermostability of a yeast mRNA, leading to estimates for ~1329 and ~2215 distinct yeast mRNAs in PARTE and DMS-seq data, respectively.

### Functional importance of genes

We used three different metrics as proxies of functional importance of genes, namely gene indispensability, mRNA expression level and evolutionary conservation. For gene indispensability, fitness measurements of 4218 single gene deletion yeast strains were downloaded from (Steinmetz, et al. 2002), and essentiality of mouse protein coding genes were extracted from (Pal, et al. 2003). Expression levels of mRNA as measured by RNA-seq in yeast and mouse were downloaded from GSE11209 (Nagalakshmi, et al. 2008) and GSE93619(Good, et al. 2017), respectively, to match the cell line/tissue used in RPL/PARIS. Evolutionary conservation is estimated inversely by the ratio between the number of nonsynonymous substitutions per non-synonymous site (dN) and the number of synonymous substitutions per synonymous site (dS) detected from one-to-one orthologs between *S. cerevisiae* and *S. bayanus* following previously describe pipelines(Zhang and Yang 2015). Briefly, orthologous proteins are identified by reciprocal best hit of BLASTp between proteomes of the two species, with the criteria of E value < 10^−20^ and alignment covering at least 80% of both orthologous sequences and at least 30-amino acid long. To avoid the influence of gene duplication, we used only one-to-one orthologous proteins. That is, we excluded any protein from a species if it is the best hit for more than one proteins in the other species. The orthologous gene pairs was re-aligned by ClustalW(Thompson, et al. 2002), filtered for gaps in alignment, and processed by PAML(Yang 2007) to calculate dN/dS.

### Evolutionary conservation of each nucleotide in yeast transcriptome

To estimate the evolutionary rate of individual sites in *S. cerevisiae* mRNA, we collected the mRNA sequences from five other post-WGD (whole-genome duplication) fungal species (*S. paradoxus*, *S. mikatae*, *S. bayanus*, *Candida glabrata*, and *S. castellii*), along with gene orthology/paralogy information among the six species from the Fungal Orthogroups Repository (Wapinski, et al. 2007). Only one-to-one orthologs in all six species were used in our analysis. We aligned orthologous mRNA sequences by ClustalW(Thompson, et al. 2002), excluding any alignment columns with gaps in any sequence. We then used GAMMA(Gu and Zhang 1997) to estimate the site-specific substitution rates of each nucleotide in each mRNA. The evolutionary conservation of a nucleotide is the inverse of its substitution rate.

### Odds ratios and MH test

We defined and computed an odds ratio (*OR*_1_) to detect within-gene correspondence between evolutionary conservation and folding specificity. To estimate the *OR*_1_, a 2 × 2 contingency table was constructed for each gene by respectively categorizing each nucleotide into one of four types on the basis of (i) whether its folding specificity is higher than the mean folding specificity of all nucleotides of the gene and (ii) whether it is more conserved than the mean level of evolutionary conservation among all nucleotides of the gene. Let the numbers of sites that fall into the four groups be: *a* (yes to both questions), *b* (yes to only question i), *c* (yes to only question ii) and *d* (no to both questions), respectively. All *a*/*b*/*c*/*d* were added by 1 as a pseudocount to avoid division by zero. We then calculated *OR*_1_ = *ad/bc*. Thus, *OR*_1_ is >1 when conserved sites of a gene tend to have high folding specificity. The function “mantelhaen.test” provided in R was used to combine *OR*_1_’s from different genes and perform the MH test (Cochran-Mantel-Haenszel chi-squared test) (see **Fig. S4**). The detection of within-gene correspondence between *T*_*m*_ and folding specificity is carried out by calculating another odds ratio (*OR*_2_) similarly, with the exception that the 2 × 2 matrix was constructed for each gene by respectively categorizing each nucleotide into one of four types on the basis of (i) whether its folding specificity is higher than the mean folding specificity of all nucleotides of the gene and (ii) whether its *T*_*m*_ is higher than mean *T*_*m*_ among all nucleotides of the gene.

### Ribosome profiling

The ribosome profiling data for yeast were obtained from GEO under accession number GSE50049(Artieri and Fraser 2014). The raw reads were quality-filtered and adaptor-trimmed before mapped onto their respective genomes. We obtained normalized ribosome density of a nucleotide by its coverage in Ribo-Seq divided by the average coverage of the transcript it belongs to. Assuming negligible ribosome drop-off, this normalized ribosome density excludes the variation in mRNA abundance and translational initiation rate, and is inversely correlated with ribosome velocity. It is noted that to exclude the influence of 5’ ribosomal “ramp”(Tuller, et al. 2010) over the detection for peak of ribosomal density, the first 200 nucleotides of every gene are removed from our analysis.

### Selective strength on point mutations affecting mRNA folding specificity

Suppose a mutation increases folding specificity and consequently increase the ribosome density on *p* codons by *q* fold. The averaged elongation speed for the mutant is *v*′ = *Lv*/(*L* − *p* + *pq*), where *v* is its original speed and *L* is the gene length in term of the number of codons. When *p* ≪ *L* and *pq* ≪ *L*, we have

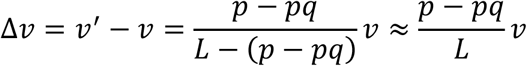

Assuming *p* = 1 codon, *q* = 1.2 (**Fig. 7A**), *L*=400 codon (average length of yeast protein) and *v* = 20 codon/s(Gilchrist and Wagner 2006; von der Haar 2008), we obtain Δ*v* = −0.01 codon/s. We then estimate the fitness effect of this Δ*v* by a previously published model(Yang, et al. 2014). Briefly, the model consists of three main components. First, by assuming a fixed number of translating ribosomes, the fitness cost of slower translational elongation is estimated by the increased time requirement for synthesizing the whole proteome for the daughter cell before cell division. Second, the quantitative relationship between elongation speed and accuracy is estimated by data from an experimental study investigating the linear trade-off between efficiency and accuracy of tRNA selection during translation (Johansson, et al. 2012). Third, the benefit of reduced translational error and/or protein misfolding is modelled by assuming a certain fraction of mistranslated proteins as misfolded, and that the misfolded proteins impose a dosage-dependent fitness cost, whose effect size is experimentally determined (Drummond and Wilke 2008). More detailed description of the model can be found in (Yang, et al. 2014).

## ACKNOWLEDGEMENT

This work was supported by the National Key R&D Program of China (grant number 2017YFA0103504 to X. C., and grant number 2018ZX10301402 to J.-R. Y.), and the start-up grant from “100 Top Talents Program” of Sun Yat-sen University (grant number 50000-18821112 to X. C. and grant number 50000-18821117 to J.-R. Y.), and by the National Natural Science Foundation of China (grant number 31671320 and 31871320 to J.-R. Y.).

## Supplementary figures and legends

**Figure S1.**
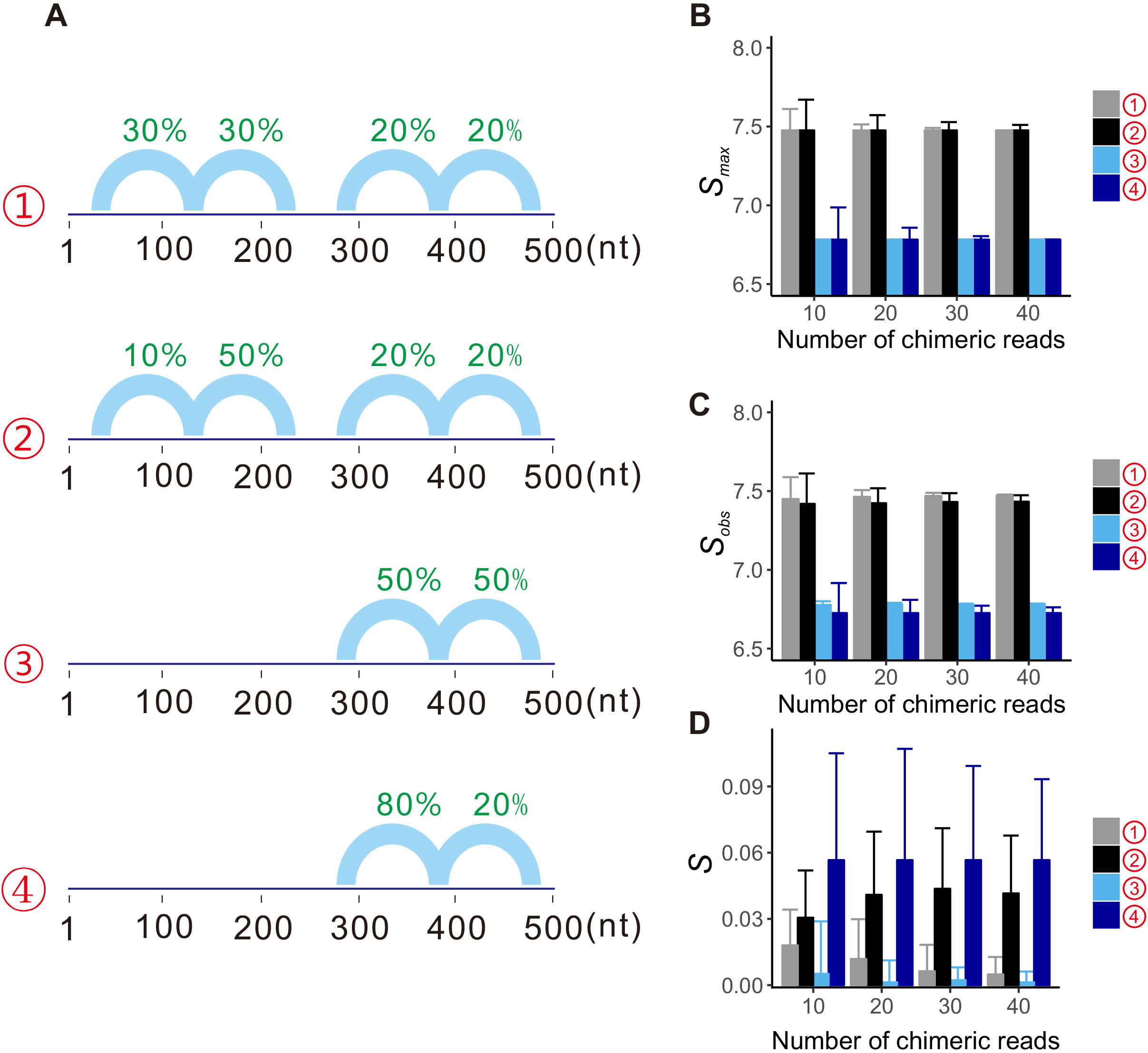
Demonstration of folding specificity calculation by an imaginary example. (A) Four hypothetical genes containing one or two regions with potential alternative folding partners were simulated. The folding partners within each gene are represented by blue arches, with green numbers indicating the probability of observing reads supporting a particular folding. For gene ➀, both regions have alternative foldings that are equally likely, i.e. the gene has no detectable signal of folding specificity (*S*=0). For gene ➁, one of the regions shown increased folding specificity since more reads (50%) support one folding, and less (10%) support the alternative folding. Gene ➂ and ➃ similarly have weak and strong signal of folding specificity, respectively, but have only one region with potential alternative foldings. To mimic the RPL/PARIS experiment, 10 to 40 chimeric reads are randomly assigned to each gene following the green probabilities, as supports for the foldings. The chimeric reads were then used to estimate *S*_max_ (B), *S*_obs_ (C) and *S* (D) for each hypothetical gene, respectively. The simulation of chimeric reads and estimation of *S*_max_, *S*_obs_ and *S* for each hypothetical gene was repeated for 1,000 times, and their mean and standard error were respectively plotted as the bar and the error bar.

**Figure S2.**
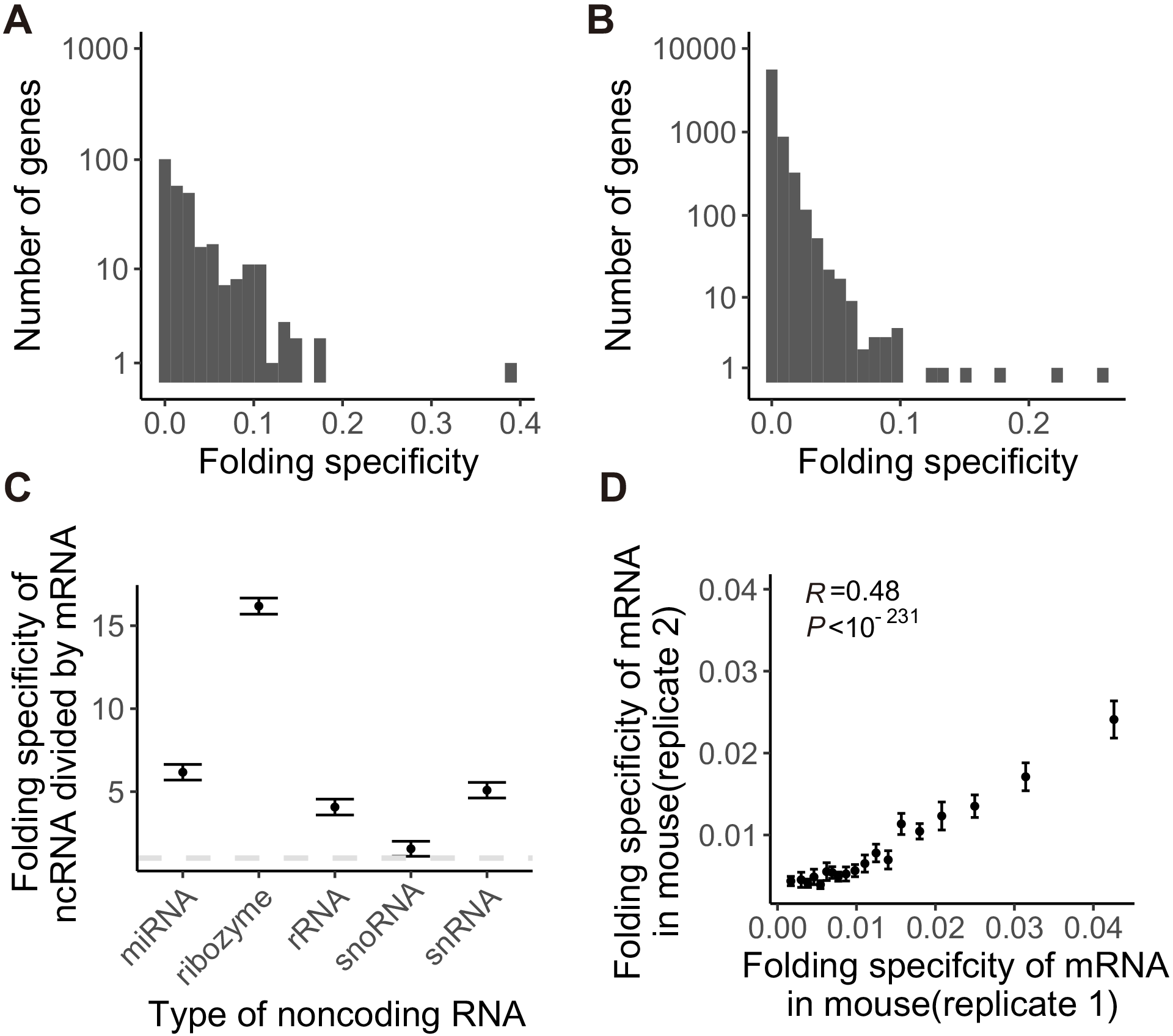
(A) Distributions of folding specificity for mRNA with at least 10 chimeric reads from RPL in yeast. (B and C) Same as Fig. 1B and C, except that folding specificity of mouse transcripts as derived from PARIS were used. Types of ncRNA presented in C include, microRNA (miRNA), ribozyme, ribosomal RNA (rRNA), small nucleolar RNA (snoRNA) and small nuclear RNA (snRNA). (D) Folding specificities of mRNA estimated from two PARIS replicates in mouse were compared. All genes with at least 5 chimeric PARIS reads were used to calculate the Pearson’s Correlation Coefficient *R* = 0.48 (*P* < 10^−231^). Genes with folding specificity > 0 were divided into 20 equal-sized groups, and the average folding specificity for each group in either PARIS replicates were plotted, where error bars indicate standard error.

**Figure S3.**
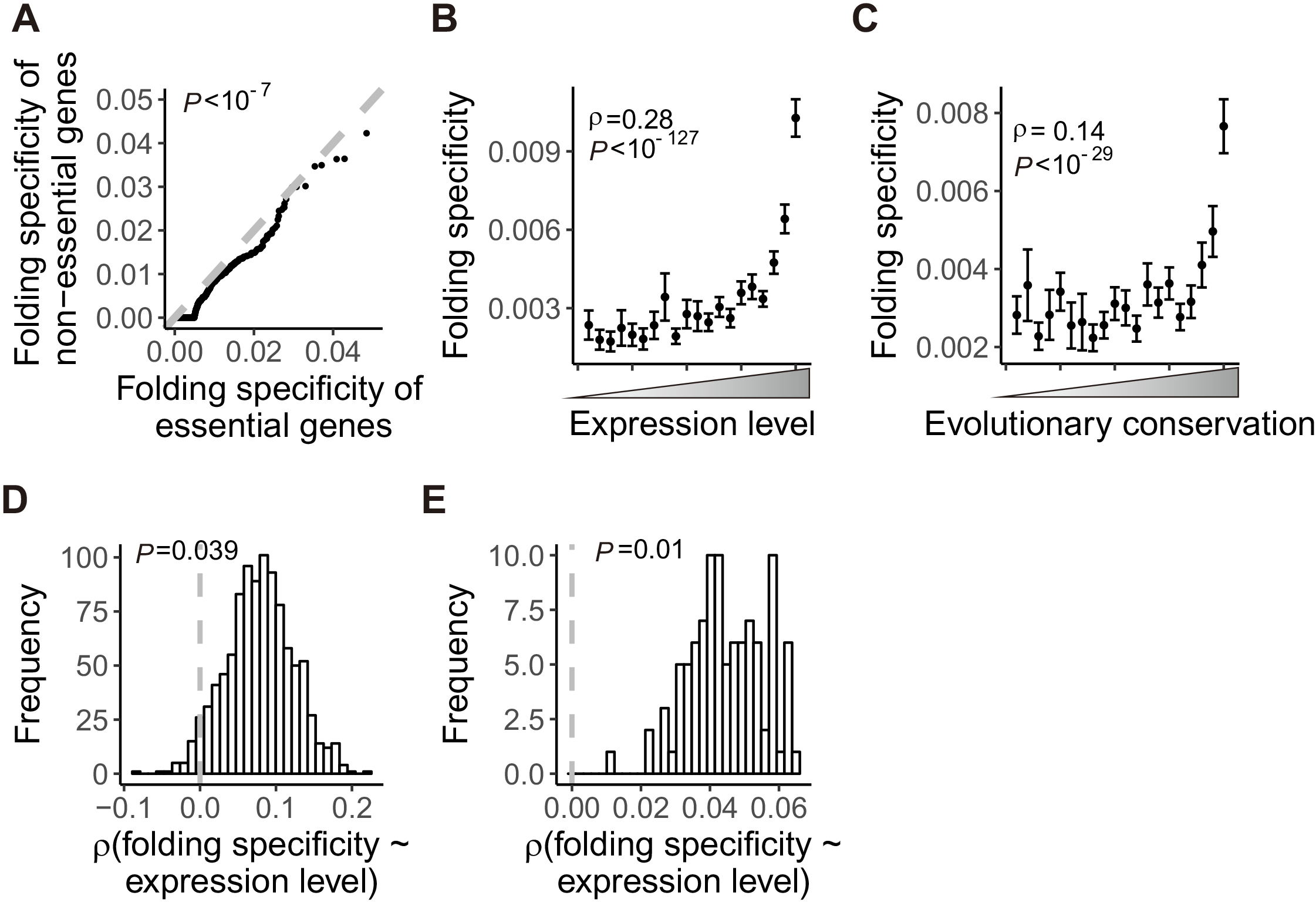
Important genes have stronger folding specificity. (A) Folding specificity of essential and nonessential protein coding genes in mouse were compared by a quantile-quantile plot. Essential genes have significantly high folding specificity than nonessential genes (*P* < 10^−7^, Wilcoxon rank sum test), as evident by the points remained below the gray line of *x* = *y*. (B and C) Same as Fig. 4B and C, except that folding specificity of mouse mRNA as derived from PARIS were used. (D and E) To exclude the possibility that stronger folding specificity in highly expressed genes is an artefact created by the abundance of chimeric reads for highly expressed mRNA, we randomly sampled five intramolecular chimeric reads from each mRNA and recalculate folding specificity. Such randomized down-sampling was repeated for 1,000 and 100 times in yeast and mouse respectively (down-sampling in mouse is too slow because of the number and length of genes), and the resulting folding specificity was still positively correlated with mRNA expression level in yeast (D) and mouse (E)

**Figure S4.**
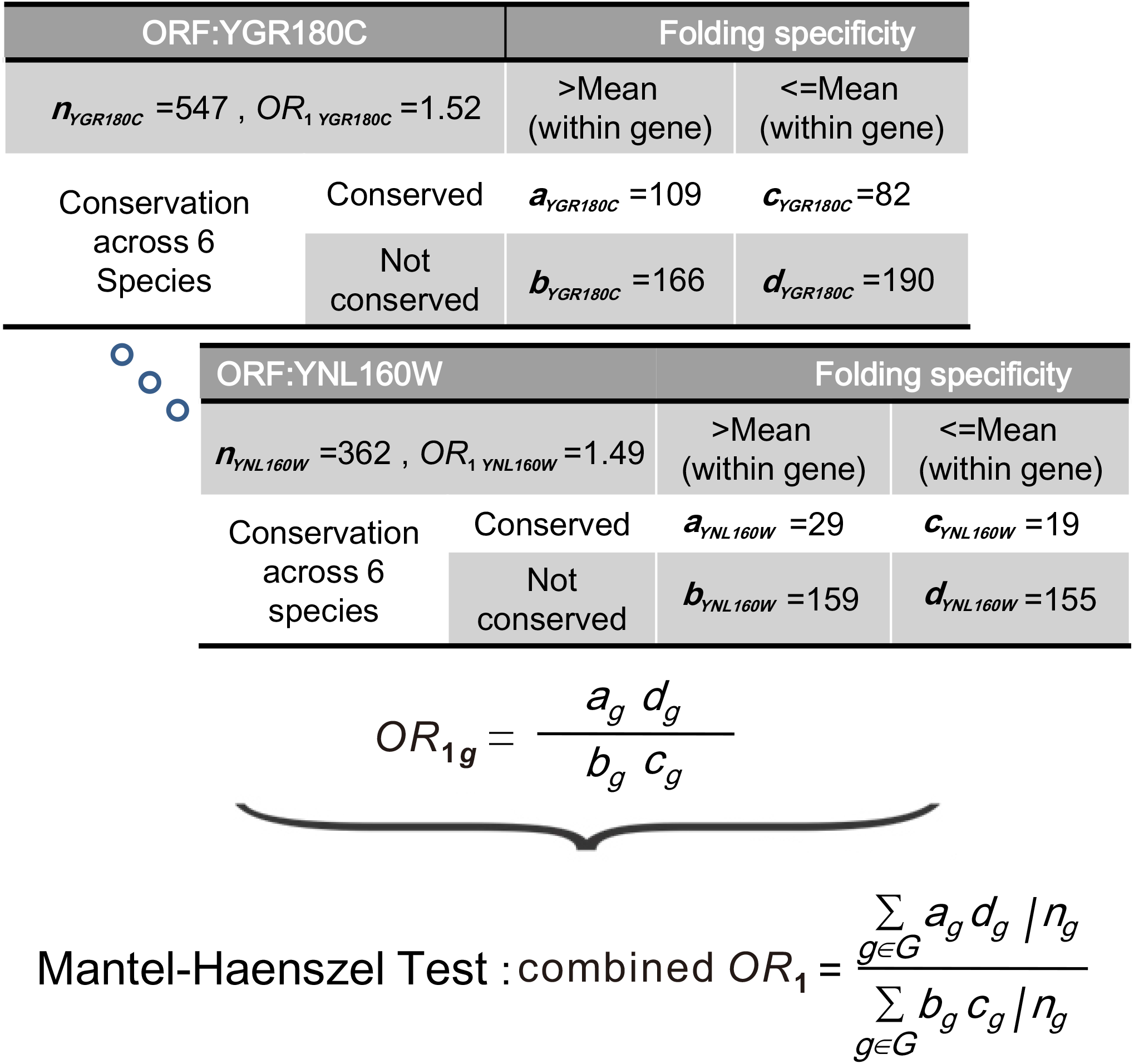
A schematic diagram of combined odds ratios calculation by the Mantel-Haenszel Procedure. A 2×2 contingency table was constructed for each gene by respectively categorizing each nucleotide into one of four types on the basis of (i) whether its folding specificity is higher than the mean folding specificity of all nucleotides of the gene and (ii) whether it is more conserved than the mean level of evolutionary conservation among all nucleotides of the gene. Let the numbers of sites that fall into the four groups be: *a* (yes to both questions), *b* (yes to only question i), *c* (yes to only question ii) and *d* (no to both questions), respectively. All *a/b/c/d* were added by 1 as a pseudocount to avoid division by zero. We then calculated *OR*_1_ = *ad/bc*. Thus, *OR*_1_ is > 1 when conserved sites of a gene tend to have high folding specificity. Then, Mantel-Haenszel Test was used for the combined *OR*_1_ calculated for all genes using the indicated formula.

**Figure S5.**
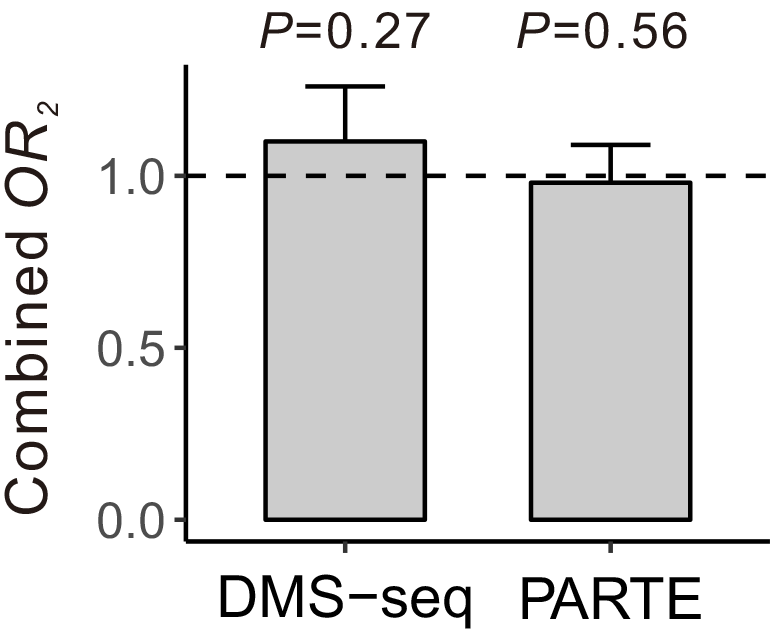
The local influence of thermodynamic stability *in vitro* on folding specificity is tested by calculation of an odds ratio (*OR*_2_) for the within-gene correspondence between folding specificity and melting temperature (*T*_*m*_) (See Materials and Methods). However, *T*_*m*_ estimated by neither DMS-seq nor PARTE display significant result, suggesting that *in vitro* thermodynamic stability has small, if any, effect on folding specificity, a result consistent with between gene analyses (Fig. 3). Combined *OR*_2_ values estimated by Mantel-Haenszel procedure are shown. Error bar indicates one standard error, estimated by bootstrapping the genes 1,000 times

**Figure S6.**
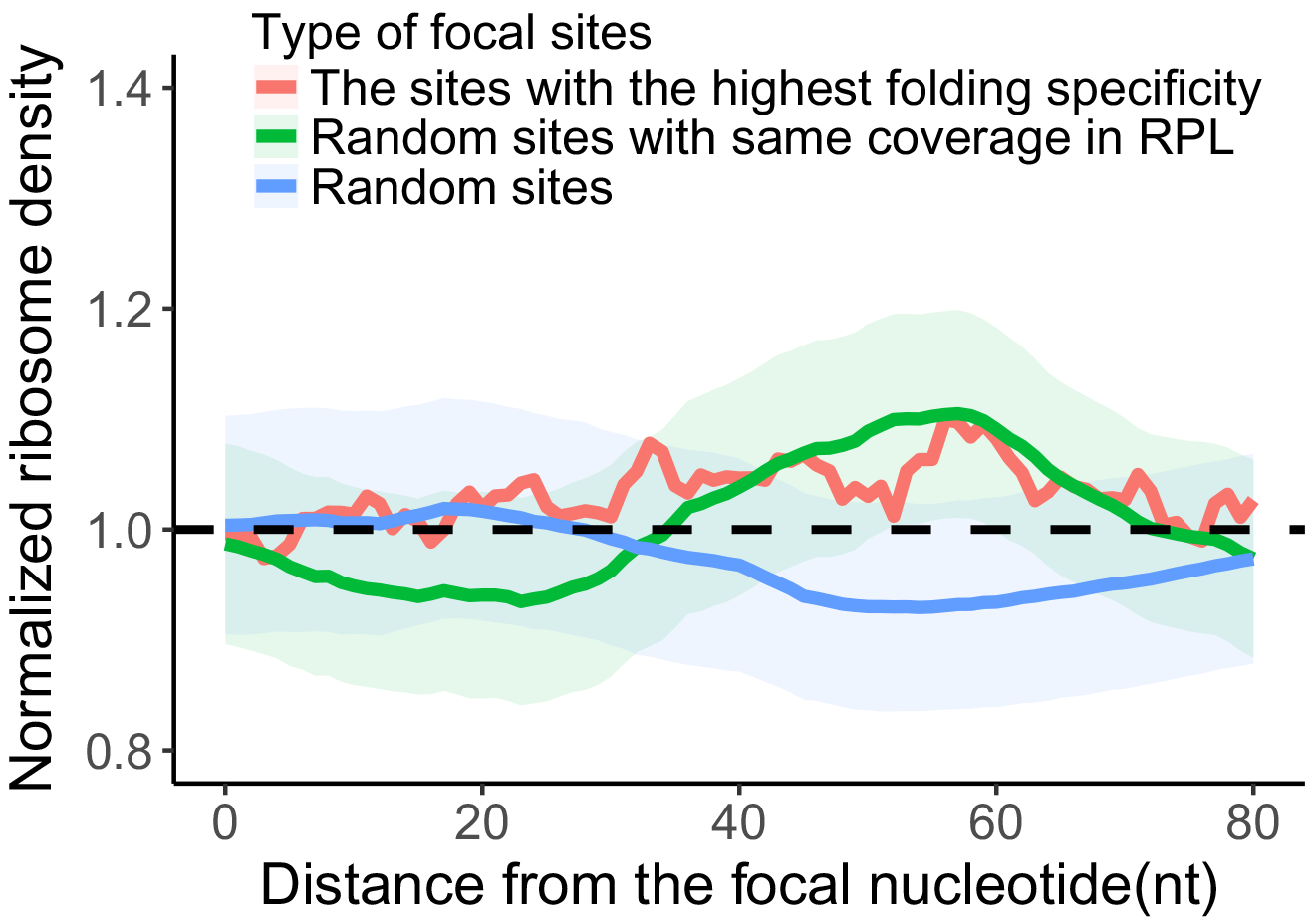
Average normalized ribosome density for the downstream of the most specifically folded nucleotides. This figure is same as that in Fig. 7A, except that the downstream of the most specifically folded (focal) site is shown.

